# Coherent Raman microscopy detects nucleolar defects through amide I peak shifts originating from β-sheets: an application to visualizing ongoing cellular senescence

**DOI:** 10.1101/2024.07.01.600896

**Authors:** Shigeo Ishibashi, Akihito Inoko, Yuki Oka, Philippe Leproux, Hideaki Kano

**Author notes:** These authors contributed equally to this work (Shigeo Ishibashi, Akihito Inoko). Correspondence should be addressed to A.I or H.K.

## Abstract

Cellular senescence occurs through the accumulation of many kinds of stresses. Senescent cells in tissues also cause various age-related disorders. Therefore, detecting them without labeling is beneficial. However, existing biomarkers have limitations of requiring fixation and labeling, or their molecular backgrounds are uncertain. Coherent anti-Stokes Raman scattering (CARS) spectroscopic imaging is a novel option because it can assess and visualize molecular structures based on their molecular fingerprint. Here, we present a new label-free method to visualize cellular senescence by obtaining molecular fingerprint signals in nucleoli using a CARS microspectroscopic system. We found the peak of the nucleolar amide I band shifted to a higher wavenumber in binuclear senescent cells, which reflects changes in the protein secondary structure from predominant α-helices to β-sheets originating from amyloid-like aggregates. Following this, we developed a procedure that can visualize the senescent cells by providing the ratios and subtractions of these two components. We also confirmed that the procedure can visualize nucleolar aggregates due to unfolded/misfolded proteins produced by proteasome inhibition. Finally, we found that this method can help visualize the nucleolar defects in naïve cells even before binucleation. Thus, our method is beneficial to evaluate ongoing cellular senescence through label-free imaging of nucleolar defects.

## Introduction

Aging in humans causes various problems such as decreases in muscle mass, bone density, and skin elasticity. These originate from senescence at the cellular level [1–3]. Cellular senescence is classically defined as the irreversible growth arrest of normal diploid cells. The accumulation of various stresses damages DNA, proteins, and lipids in cells, thereby impairing cellular homeostasis, which causes cellular senescence [4–7]. Hence, cellular senescence also plays a role in preventing the onset of cancer. However, cellular senescence also causes the depletion of stem cells and the release of cytokines from aged cells, which results in the age-related disorders mentioned above [2,4]. Therefore, detecting and evaluating senescent cells is crucial for research on age-related disorders and their prevention [8–11].

Cellular senescence has long been evaluated in chemically fixed and labeled cells [12,13]. These include empirical detection of senescence-associated β-galactosidase (SA-β-gal) and immunostaining for mechanistically assured biomarkers, such as p16, p21, Rb, and p53. In contrast, accurate methods for detecting unlabeled senescent cells are few [13–16]. The aggregation and enlargement of nucleoli are major indicators of cellular senescence under an optical microscope [12,14–16]. However, the observation is limited to cells in cultured dishes, and the assessment of molecular properties from the images is difficult. Developing a method for detecting unlabeled senescent cells based on the accurate molecular background is desired.

Optimization of Raman microscopy is suitable for this purpose. Spontaneous Raman scattering (SpR) microscopy is a candidate to identify senescent cells [17–21]. However, the molecular basis of this technique is not well understood. Coherent Raman scattering (CRS) microscopy has higher sensitivity than SpR microscopy and is more suitable for this purpose. It utilizes the nonlinear optical process generated by pulsed light with high peak intensity [22–30]. In particular, ultra-multiplex coherent anti-Stokes Raman scattering (CARS) microscopy, “a subtype” of CRS microscopy that we developed, is superior to other CRS microscopies because it enables the extraction of detailed molecular information encoded in fingerprint CARS spectra. Thus, CARS microspectroscopy may be beneficial for label-free imaging of senescent cells based on its spectroscopic signature originating from its molecular fingerprint.

The nucleolus is the candidate organelle to determine cellular senescence. Its basic function is to produce ribosomes, an apparatus that synthesizes proteins [31]. Notably, recent research has highlighted their function in maintaining the quality of proteins particularly under conditions of stress. A nucleolus disorder is likely to be associated with cellular dormancy [1,32,33] and cellular senescence [34]. Stress-induced misfolded/unfolded proteins are stored in nucleoli for refolding during cell recovery. However, prolonged stresses make the stored proteins more solid, which results in cell dormancy [35]. Accumulation of nucleolar protein is also observed in senescent cells [34], with some of them identified as amyloid aggregates [36]. Overall, accumulation and aggregation of nucleolar protein are a characteristic of cells that are exposed to stresses and undergo senescence.

The nucleolus is a membrane-less organelle whose structural changes link to its physiological function and pathology. The nucleolus contains various ribosomal proteins and rRNA, whose architectural changes uniquely correlate with the function and occur according to liquid-liquid phase separation (LLPS) [37,38]. LLPS is a structural phase transition mainly caused by intrinsically disordered proteins with low-complexity domains that potentially form β-sheets [39]. These proteins form liquid droplets that possibly contain cross-β polymers, cross assembled β-sheets [40]. They become more rigid as gels and aggregates as the progression of phase separation [40–43]. One of the important physiological functions of LLPS in membrane-less organelles is the spatial regulation of translation and transcription by binding to nucleic acids such as RNA and DNA, as nucleoli show, that is summarized as the maintenance of protein quality control, in other words, proteostasis.

In contrast, the abnormal aggregations of the β-sheets are called “amyloids,” pathological intracellular and extracellular structures. During amyloidization, proteins undergo various phases such as unfolding, nucleation, elongation, and stationary [41,43,44]. However, all these phases involve β-sheet formation due to protein misfolding. Amyloid aggregates have been characterized spectroscopically in patients with systemic amyloidosis, ALS, Alzheimer’s disease, and prion disease, as well as ones produced *in vitro.* For example, Raman spectra of amyloid aggregates *in vitro* show a higher shift of amide I peak wavenumber than that of corresponding monomers, which was confirmed by insulin [45,46], lysozyme [47], and amyloid beta [48]. However, these aggregates are large, and there is little spectroscopic information available about intracellular micro amyloid. In particular, attempts to focus on cellular senescence and apply it to visualization have not been developed, because of the limitations of biological materials and the difficulties in cross-disciplinary experiments.

In this study, we observed binuclear senescent cells with CARS microscopy and found the peak of the nucleolar amide I band shifted to a higher wavenumber in the spectra. This shift reflects changes in the amount of protein secondary structure from α-helices to β-sheets originating from amyloid-like aggregates. We also developed a procedure that can visualize such nucleolar defects by analyzing the ratio and subtraction of these two components. Finally, the procedure could visualize nucleolar defects in naïve cells before binucleation. Thus, our method is beneficial to evaluate ongoing cellular senescence through imaging their nucleolar defects without labeling.

## Results

### Drug-induced binuclear senescent cells show higher-wavenumber peak shifts of amide I in nucleoli originating from β-sheets

We used human primary cells (prostate epithelial cells, PrECs) to evaluate cellular senescence because they are not cancer cells but normal diploids that can senesce. We treated them with dihydrocytochalasin B (DCB), an inhibitor of actin polymerization, because DCB stably makes cells polyploidy and can induce cellular senescence [49] (Fig. 1).

**Figure 1.**
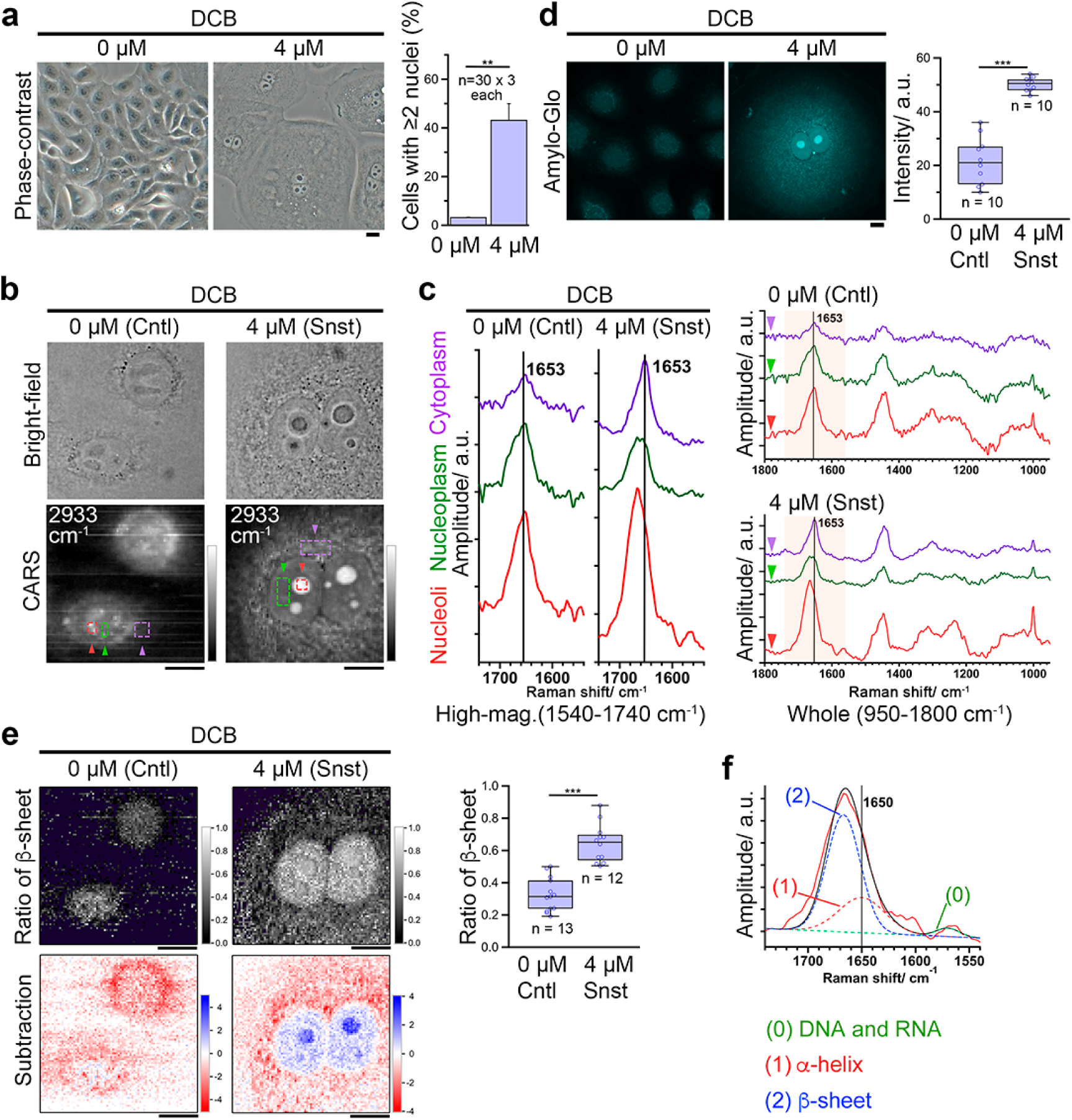
Drug-induced binuclear senescent cells show nucleolar amide I higher-wavenumber peak shifts originating from β-sheets. PrECs were treated with or without 4 µM DCB for 72 h and analyzed by the following microscopies. **(a)** Left, phase-contrast images confirm the effective induction of polyploid cells with ≥ 2 nuclei through DCB treatment. Right, the quantification. n = 3 (30 counts each). Hereafter, cells are defined as follows: ‘Control (Cntl),’ cells in 0 µM and with a single nucleus (<118 µm^2^); ‘Senescent (Snst),’ cells in 4 µM and with double nuclei. **(b)** Bright-field images (upper) and CARS images (lower) at wavenumber 2933 cm^-1^ (Im[χ^(3)^] spectral images). **(c)** Raman spectra (Im[χ^(3)^] spectra) of cytoplasm (purple), nucleoplasm (green), and nucleoli (red), whose origin is indicated in **b** by arrowhead and dotted rectangles (averaged). Left, higher magnifications of amide I vibration band. The peak shift was shown in comparison to the peak value of cytoplasm at 1653 cm^-1^ (vertical line). Right, whole spectra, where magnified areas were colored. **(d)** Left, fluorescent images with Amylo-Glo staining, an amyloid indicator dye. Right, the quantification of emission intensity in the nucleoli (20 × 20 pixels). **(e)** Ratiometric and subtraction images based on Gaussian function fitting of the protein amide I vibration band. Upper, images of 𝑔_2, 𝕩_⁄(𝑔_1, 𝕩_ + 𝑔_2, 𝕩_). Lower, images of 𝑔_2, 𝕩_ − 𝑔_1, 𝕩_. Right, quantification of the β-sheet by 𝑔_2, 𝕩_⁄(𝑔_1, 𝕩_ + 𝑔_2, 𝕩_). A cross-sectional average of the largest nucleolus in each nucleus was used. **(f)** An example of the amide I vibration band (1540–1740 cm^-1^) of the nucleolar Raman spectrum fitted using three Gaussian functions. See also the Methods section. \*\**p* < 0.01, \*\*\**p* < 0.001. Bars, 10 µm.

At first, we confirmed the formation of adequate binucleated cells, a hallmark of cellular senescence, in the DCB-treated population (Fig 1a). Then, we compared the binuclear senescent cells with control cells. Hereafter, we defined control cells as mononuclear cells without treatment and with nuclei smaller than a certain standard (cross-sectional area of 118 μm^2^) because cells with larger nuclei are potentially polyploids that means in senescence. We used multiple types of microscopies including phase-contrast, bright-field, and CARS microscopy, and confirmed that binucleate senescent cells have larger nuclei and nucleoli than control cells (Fig. 1a,b).

Secondly, we performed examinations with our CARS microscopy (Fig. 1b,c). Notably, we found that the senescent cells exhibited the characteristic features of region-averaged Im[χ^(3)^] CARS spectra regarding the peak positions of the amide I band in nucleoli and nucleoplasm (Fig. 1b,c; red and green, respectively). These are: (1) the peak wavenumbers were higher than those of the control cells, and (2) these peak wavenumbers were also higher than those in the cytoplasm of the same cell (typically at 1653 cm^-1^). In detail, the peak shifts in the nucleoplasm were not distinct compared to those in the nucleoli.

Thirdly, we tried to clarify the origin of the high-wavenumber peak shift of amide I. We speculated that the peak shift originated from the accumulated amyloid-like aggregates, as amyloid aggregates *in vitro* show similar peak shifts [45–48], and stress-induced nucleolar aggregates contain similar misfolded proteins [32,33,50,51]. We, therefore, stained cells with an amyloid aggregate indicator dye, Amylo-Glo [52]. We found that the nucleoli of the binucleated senescent cells showed a significantly brighter fluorescence signal than that of untreated cells (Fig. 1d), confirming the presence of nucleolar amyloid-like aggregates. This implies that the β-sheets originating from misfolded/unfolded amyloid protein accumulate [33,35], thereby causing the amide I peak shift because amide I of β-sheets has a peak at a higher wavenumber than that of α-helices [45,48,53].

Therefore, we finally tried to visualize senescent cells by utilizing the nucleolar amide I peak shift. We developed a procedure and reconstructed spectroscopic images as follows (Fig. 1e,f). First, we fit the amide I band using three Gaussian functions with the following different peak wavenumbers, that we assigned to: (0) purine ring skeleton of the DNA and RNA (peak wavenumber: 1570 cm^-1^), (1) amide I of the protein ‘α-helix structure’ (peak wavenumber: 1650 cm^-1^) and cis-type carbon double bonds of unsaturated lipids, and (2) amide I of the protein ‘β-sheet structure’ (peak wavenumber: 1667 cm^-1^) (Fig. 1f, see also the Methods section). Then, we used (1) and (2) (i.e., ‘α-helices’ and ‘β-sheets’) and reconstructed the ratiometric images and subtraction ones that are calculated using 𝑔_2, 𝕩_/(𝑔_1, 𝕩_ + 𝑔_2, 𝕩_) and 𝑔_2, 𝕩_ − 𝑔_1, 𝕩_, respectively. Here, 𝑔_i, 𝕩_ represents the integrated band intensity for (1) and (2) at a particular spatial point 𝕏.

Using this procedure, we firstly estimated the amount of nucleolar amyloid aggregate through 𝑔_2, 𝕩_/(𝑔_1, 𝕩_ + 𝑔_2, 𝕩_), the ratio indicating the occupancy of β-sheets. We compared them at the largest nucleoli in each nucleus and detected higher values in DCB-treated senescent cells (Fig. 1e, right graph). However, the ratiometric images also showed higher values in the nucleoplasm (Fig. 1e, upper panel), which differed from Amylo-Glo staining (Fig. 1d). Secondly, we estimated the amount of nucleolar amyloid aggregate through 𝑔_2, 𝕩_ − 𝑔_1, 𝕩_, the subtraction indicating the relative amount of β-sheet. Notably, the subtraction image clearly visualized the nucleoli in the senescent cells (Fig. 1e, lower panel), demonstrating the accumulation of β-sheets in nucleoli rather than the nucleoplasm, which is similar to Amylo-Glo staining (Fig. 1d).

Thus, our CARS microscopy presents a new procedure for evaluating senescence through the detection of nucleolar amyloid-like aggregates, whose technical advantages are (1) allowing the quantification of protein secondary structures and (2) “not requiring a label”.

### Naturally binucleated senescent cells also show nucleolar amide I higher-wavenumber peak shifts originating from β-sheets

Next, we tried to assess naturally occurring senescent cells with our CARS microscopy method (Fig. 2). We noticed naturally binucleated senescent cells occasionally occurring in naïve PrECs (arrowheads in Fig. 2a). We compared them with control cells and confirmed that the nucleolus of naturally binucleated senescent cells was larger than that of control cells (Fig. 2b, the bright-field and CARS images (Im[χ^(3)^]) at 2933 cm^-1^). Then, we compared their region-averaged Im[χ^(3)^] spectra. As expected, the peak position of the amide I band in nucleoli and nucleoplasm exhibited similar peak shifts as those observed in DCB-treated cells (Fig. 2c). The presence of nucleolar amyloid-like aggregates was also confirmed by Amylo-Glo (Fig. 2d).

**Figure 2.**
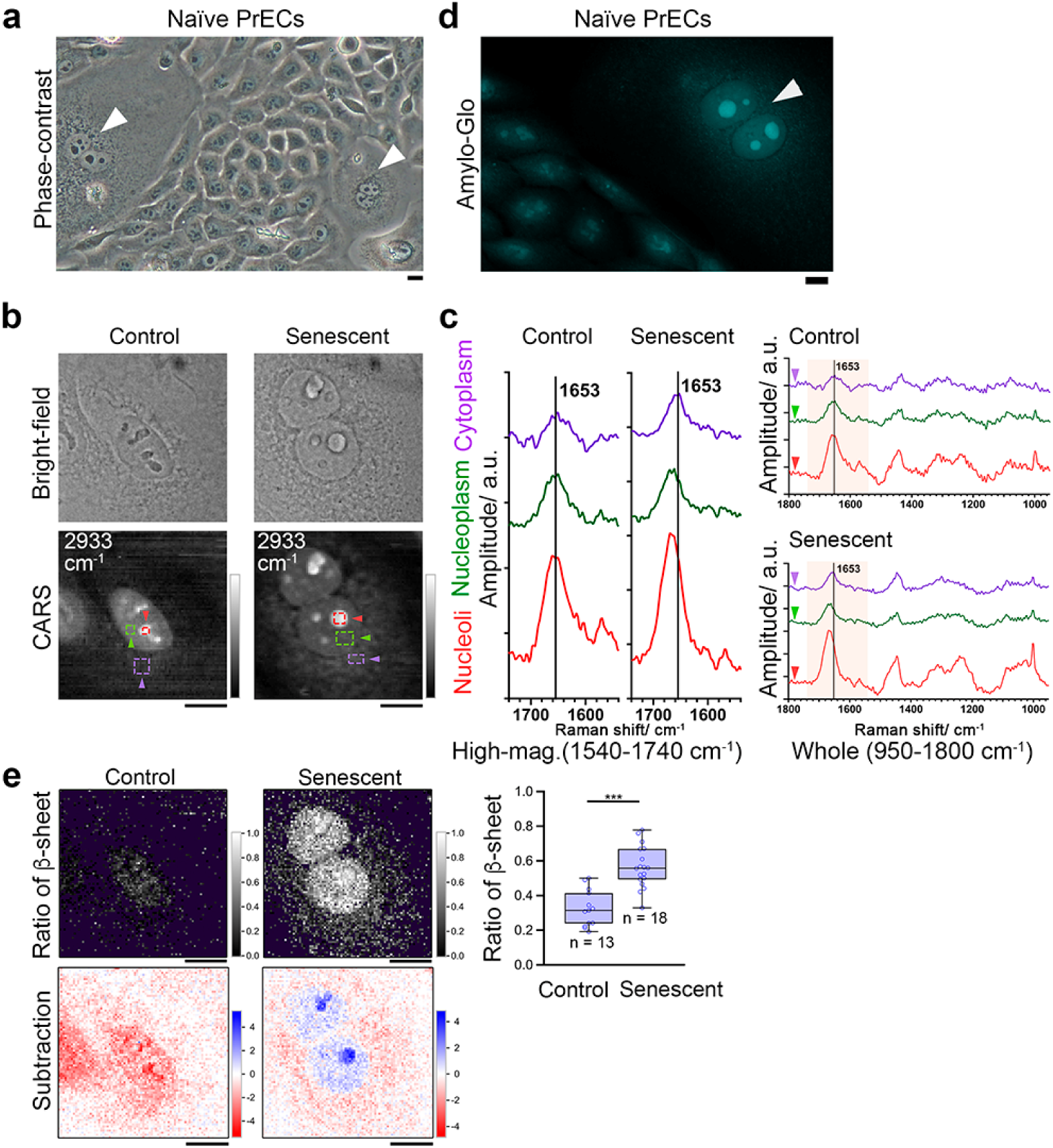
Naturally binucleated senescent cells also show nucleolar amide I higher-wavenumber peak shifts originating from β-sheets. Naturally binucleated senescent PrECs were analyzed by following microscopes. **(a)** Phase-contrast images showing naturally binucleated senescent cells (arrowheads). **(b)** Bright-field images and CARS images at wavenumber 2933 cm^-1^ (Im[χ^(3)^] spectral images). **(c)** The fingerprint Raman spectra averaged in each of the arrowhead-marked regions, in the same manner as Fig. 1. Left, higher magnifications of amide I vibration band. **(d)** Fluorescent images with Amylo-Glo staining, an amyloid indicator dye. **(e)** Ratiometric and subtraction images in the same manner as Fig. 1. \*\*\**p* < 0.001. Bars, 10 µm.

We again performed the amide I band-based ratiometric and subtraction image analysis with them and examined their validity as indicators for natural senescent cells. Our findings revealed that the nucleoli of the natural senescent cells also contain more proteins with β-sheets (Fig. 2e; right graph, upper panel). The subtraction image demonstrates that the nucleoli of the natural senescent cells also contain more proteins with β-sheets than the nucleoplasm of the cells (Fig. 2e, lower panel). Overall, we confirmed that our proposed senescent cell indicators are also effective for the detection of natural senescent cells.

### Proteasome inhibition also causes nucleolar amide I higher-wavenumber peak shifts originating from unfolded/misfolded protein aggregates

Nucleolar amyloid aggregates are also produced through proteasome inhibition that increases unfolded proteins [32]. As such, we treated PrECs with a proteosome inhibitor MG132 and optically analyzed them in the same manner as described above (Fig. 3). We confirmed that the proteasome inhibition caused the aggregation of nucleoli in the phase-contrast image (Fig. 3a). Notably, our CARS microscopy revealed that the amide I band of the MG132-treated cells exhibited almost the same peak shifts in nucleoli and nucleoplasm, as chemically-induced and naturally occurring senescent cells (Fig. 3b,c). The presence of amyloid-like aggregates was confirmed by Amylo-Glo staining as in the case of the two types of senescent cells (Fig. 3d).

**Figure 3.**
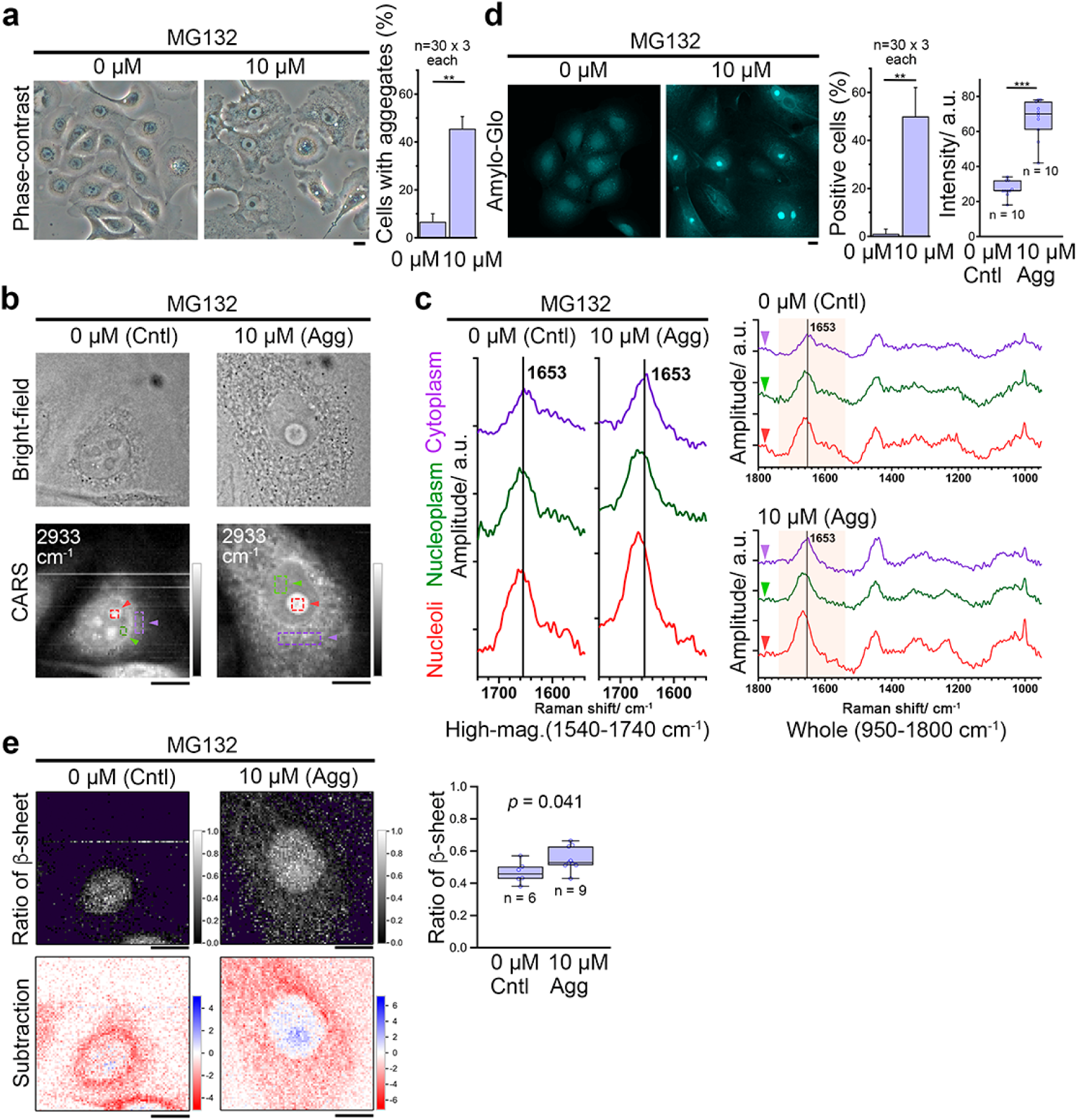
Proteasome inhibition also causes nucleolar amide I higher-wavenumber peak shifts originating from unfolded/misfolded protein aggregates. PrECs were treated with or without 10 µM MG132 for 8 h and analyzed by following microscopies. **(a)** Left, phase-contrast images confirm the formation of aggregates through MG132 treatment. Right, the quantification. 30 cells were counted in each group (n = 3). Hereafter, cells are defined as follows: ‘Control (Cntl),’ cells in 0 µM and with mononuclei (<118 µm^2^); ‘ Aggregate (Agg),’ cells in 10 µM and with aggregates. **(b)** Bright-field images of the cells (upper) and CARS images (lower) at wavenumber 2933 cm^-1^ (Im[χ^(3)^] spectral images). **(c)** The fingerprint Raman spectra averaged in each of the arrowhead-marked regions in the same manner as Fig. 1. **(d)** Left, fluorescent images with Amylo-Glo staining, an amyloid indicator dye. Middle, percentages of Amylo-Glo-positive cells. n = 3 (30 counts each). Right, intensity of the spots in the nucleoli (20 × 20 pixels); n = 10. **(e)** Ratiometric and subtraction images in the same manner as Fig. 1. \*\**p* < 0.01,\*\*\**p* < 0.001. Bars, 10 µm.

We again performed the amide I band-based ratiometric and subtraction image analysis (Fig. 3e). They revealed that the nucleoli of the MG132-treated cells also contain more β-sheets (Fig. 3e; right graph, upper panel). The subtraction image demonstrates that the nucleoli of MG132-treated cells also contain more proteins with β-sheets than the nucleoplasm of the cells (Fig. 3e, lower). Thus, our methods can visualize increased nucleolar defects with β-sheets originating from misfolded/unfolded protein-derived amyloids regardless of the causes.

### CARS detects ongoing cellular senescence through nucleolar defects

The size of the nucleolus is considered to correlate with the degree of cellular senescence [14]. As such, we finally evaluated the correlation between the nucleolus size and the CARS Im[χ^(3)^] spectra in mononuclear cells (Fig. 4).

**Figure 4.**
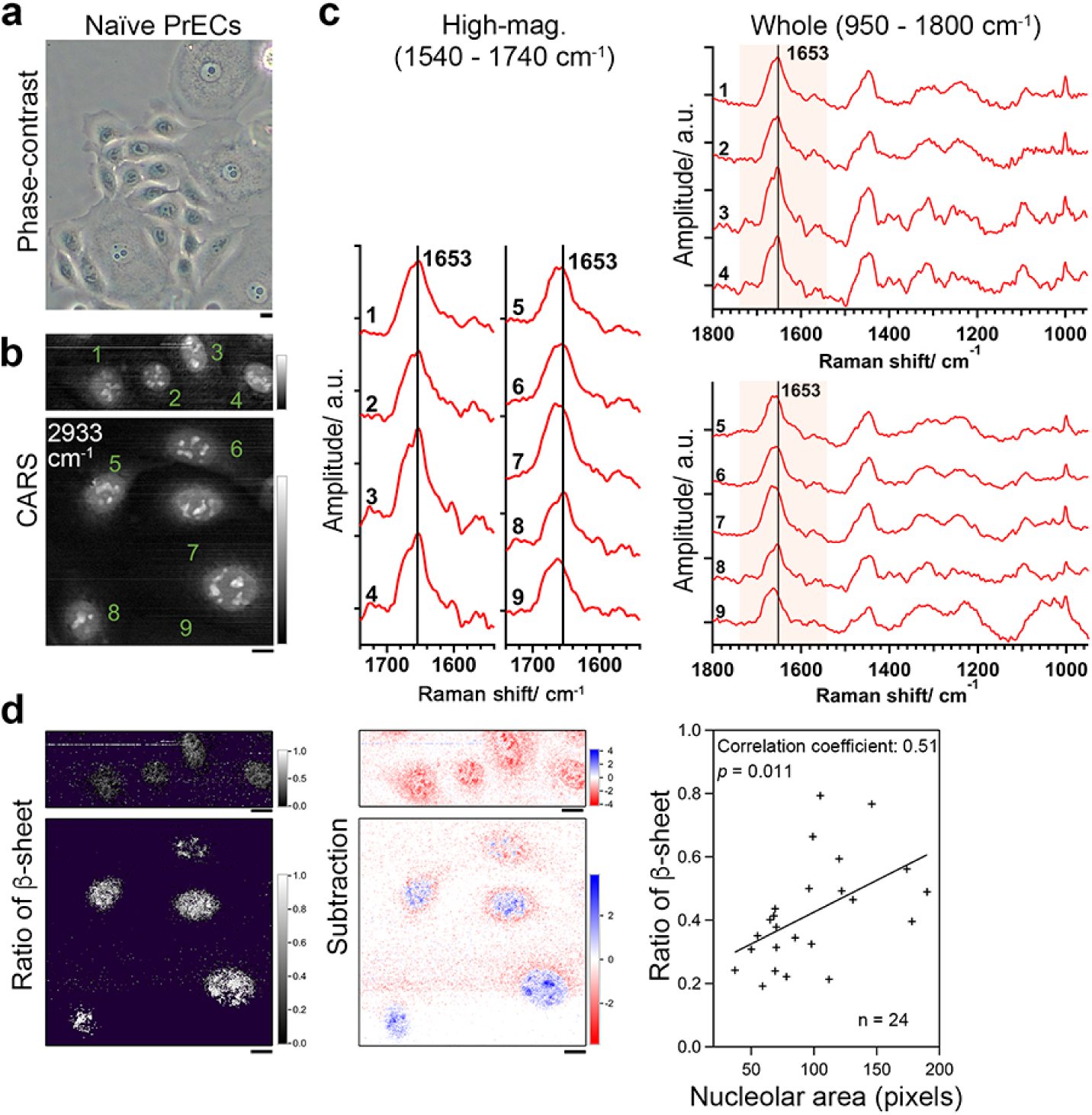
CARS detects ongoing cellular senescence through nucleolar defects. Measurements of naïve mononuclear PrECs with various nucleolar sizes with microscopes. **(a)** Phase-contrast images show a variety of nucleolar sizes among mononucleated cells. **(b)** CARS image at wavenumber 2933 cm^-1^ (Im[χ^(3)^] spectral images), confirming the variation of nucleolar size. The numbers correspond to the spectra of (c). **(c)** The fingerprint Raman spectra of the largest nucleolus in each nucleus. Each spectrum is average in the corresponding nucleolus. Left, higher magnifications. Right, whole spectra. Vertical lines at 1653 cm^-1^ indicate the peak in the protein amide I vibrational band of Ctrl nucleoli. **(d)** Left and middle, ratiometric and subtraction images in the same manner as Fig. 1. Right, the cross-sectional area of the largest nucleolus in each cell (as a counted number of pixels) vs. the ratio of β-sheet (𝑔_2, 𝕩_/(𝑔_1, 𝕩_ + 𝑔_2, 𝕩_)). The correlation coefficients and *p*-values shown in the graph are convincing. Bars, 10 µm.

Naïve mononuclear PrECs showed considerable variations in the sizes of the cells and nucleoli (Fig. 4a, phase-contrast image; Fig. 4b, CARS (Im[χ^(3)^]) images at 2933 cm^-1^, representative nine cells within total 24 cells). Notably, a comparison of the average Im[χ^(3)^] spectra of the nucleolar region among the cells suggests the variation in the amide I peak shift (Fig. 4c). The ratiometric and subtraction images revealed that the occupancy ratio and the amount of the β-sheet component tended to increase with the size of the nucleoli (Fig. 4d, left and middle panel). The cross-sectional averaged values of 𝑔_2, 𝕩_/(𝑔_1, 𝕩_ + 𝑔_2, 𝕩_) for the size of the largest nucleolus in each mononuclear cell are shown as a scatterplot (Fig. 4d, right graph), that result in the linear regression confirming the significant correlation (*p* = 0.011). Thus, our procedure can detect unfolded protein accumulating into nucleolar aggregates even in mononuclear cells, which enables the evaluation of ongoing cellular senescence through nucleolar unfolded proteins before binucleation.

## Discussion

In this study, we developed new detection criteria for the label-free senescent cell discrimination method using ultra-multiplex CARS microspectroscopy (Fig. 1 and 2). We found a high-wavenumber shift of the amide I peak in the amyloid-like nucleolar aggregates in senescent cells, whose volumes are relatively smaller than pathological amyloids. We further found the increase in proteinous β-sheet components in the amide I band through Gaussian function fitting. Conclusively, we established visible senescence indicators as the ratios of the spectral components for the α-helix and β-sheets and their corresponding subtractions. These features of nucleoli are also common in proteasome inhibitor-treated cells (Fig. 3) and pre-binuclear ongoing senescent cells (Fig. 4). This is due to the large number of β-sheets that are thought to originate from unfolded proteins. To our knowledge, this study is the first demonstration that the CARS microscopy can identify nucleolar small amyloid aggregates-specific spectra, that can advantageously be applied to the detection of senescent cells. SpR microscopy has already been used to examine senescent cells. However, the underlying molecular features are unclear, and the nucleolar amide I peak shift has not been previously reported [17–21]. The ability to detect another type of amyloid, “amyloid plaque in Alzheimer’s disease brain,” was previously compared between SpR microscopy and stimulated Raman scattering (SRS) microscopy, another subtype of CRS microscopy [54]. Both microscopies showed Raman spectra with a peak shift of the amide I band to a higher wavenumber than that of other tissues. However, SpR microscopy was affected by autofluorescence in the tissues. Acquisition of the expected spectrum was complicated, due to analyses of the three-dimensional information consisting of Raman spectra at each two-dimensional coordinate point [54]. Overall, CRS microscopy is more precise in positional resolution than SpR microscopy, but the detection of senescent cells by CRS microscopy has not been reported until our report.

One reason is that our CARS microscopy is advantageous in combination with spectroscopy compared to SRS microscopy. Among the reported CRS microscopes, our ultra-multiplex CARS microscopy used in this study has high sensitivity and the capability of simultaneous detection with ultrabroad spectral coverage [23].

Our CARS microscopy could focus on the general secondary structure of proteins in nucleoli. This generality is an advantage of CARS microscopy compared to conventional molecular analysis with protein sequence-specific antibodies. Indeed, nucleoli are composed of diverse ribosomal proteins and interact with various proteins such as VHL and MDM2 [33,50]. The conventional methods of focusing on individual proteins and tracking them may be effective for the analysis of each protein. However, it may overlook changes in the whole nucleolus. In contrast, our CARS microscope has the potential to enable the measurement of whole-cell phenomena by focusing on protein structures. In the future, it will be a powerful tool for “phasing biology,” a new biology focusing on protein structure during LLPS [42,55].

In this study, we could detect nucleolar amyloid formation possibly at early stages (Fig. 4). However, currently we have not clarified what stage of β-sheets we detected. Amyloid formation progresses from looser states to gels and aggregations [34,42]. It would be beneficial to consider other schemes involving oligomers and protofibrils [43]. Focusing on each stage and comprehensively evaluating the differences in CARS spectra will be addressed in future studies, which will be beneficial for phasing biology.

Our observation indicated that nucleolar CARS values are likely less affected by proteasome inhibitors than by cellular senescence, suggesting that the origin of the β-sheets detected is merely the aggregation of misfolded proteins, and the effect of conjugated ubiquitin by inhibiting degradation is small. However, the components and manners of intranuclear aggregates are indeed diverse [56]. It is also modified by chaperone proteins such as HSP70 [35]. Furthermore, the accumulation of substances in the nucleolus differs depending on stress, such as VHL accumulation in hypoxia. Conversely, some ribosomal proteins are released, when nucleolar stress occurs due to impaired ribosomal RNA synthesis [34]. Thus, focusing on each stress and comprehensively examining the differences in nucleolar CARS spectra will further enhance the applicability of this visualization technique to cell biology.

In conclusion, this study focused on the nucleolus of senescent cells through our CARS microscopy method, where a tiny accumulation of low-quality proteins could be visualized without labeling. However, the kind of accumulating protein and the place of accumulation varies depending on the phenomenon [57]. In the future, it will be important to focus on protein quality control performed in different situations and locations such as mitochondrial stress and autophagy. Evaluating dynamic cell alteration such as cell differentiation and cancer is also valuable. As such, collecting characteristic CARS spectra from further biological models will be important to expand the application of our method.

## Methods

### Preparation of cells

PrECs were purchased from Lonza (Basel, Switzerland) and expanded for long-term culture under feeder-free conditions, according to a previous report [58]. The manufacturer confirmed that the cells were isolated from donated human tissue after obtaining permission for their use in research applications by informed consent or legal authorization. No further ethical approval was required for this study since we used commercially provided human cells following the internal review of our institutional review board.

The culture medium comprised 10 µM Y-27632 (1000583, Cayman Chemical, Ann Arbor, MI, USA), 1 µM DMH1 (041-33881, Wako, Tokyo, Japan), and 1 µM A-83-01 (039-24111, Wako) [58], all mixed in F-medium [59] containing 3:1 (v/v) Ham’s F-12 (087-08335, Wako) and Dulbecco’s modified Eagle medium (043-30085, Wako), supplemented with 5% fetal bovine serum, 0.4 µg/mL hydrocortisone (086-10191, Wako), 5 µg/mL insulin (I9278, Sigma, St. Louis, MO, USA), 8.4 ng/mL cholera toxin (030-20621, Wako), 10 ng/mL epidermal growth factor (059-07873, Wako), 24 µg/mL adenine (A2786, Wako), 100 µg/mL streptomycin (Meiji Seika, Tokyo, Japan), 100 U/mL penicillin (Meiji Seika), 250 ng/mL amphotericin B (A2942, Sigma), and 10 µg/mL gentamicin (078-06061, Wako). As necessary, the cells were treated with 4 µM DCB (250225, Merck, Darmstadt, Germany) for 72 h or 10 µM MG132 (M8699, Sigma) for 8 h. All cells were maintained at 37°C in a humidified incubator with 5% CO_2_ and used under proliferation conditions.

For microscopic observations, the cells were grown on coverslips (Iwaki Glass Co., Ltd., Tokyo, Japan) and fixed with 1% formaldehyde in phosphate-buffered saline (PBS) for 15 min. After washing thrice in PBS, the cells were observed directly or following treatment with Amylo-Glo® (TR-300-AG, Biosensis, SA, Australia) according to the manufacturer’s protocol. All the samples were immediately observed. The indicator dye Amylo-Glo increases the emission intensity at a wavelength of approximately 440 nm by binding to amyloid aggregates [52].

For CARS microscopy without staining, the cells were grown on coverslips (Iwaki Glass) and fixed with 1% formaldehyde in PBS for 15 min. After washing thrice with PBS, the cells were maintained in PBS with a ProClin preservative. After fixation, the cells were stored in liquid form at 10°C or less until CARS microscopy measurements were performed. Before the measurement, a cover glass was placed on the glass slide and sealed with a topcoat to prevent it from drying.

Control cells were defined as mononuclear cells with nuclei smaller than a certain standard (cross-sectional area of 118 μm^2^). All measurements were performed after fixation.

### Conventional microscopy image acquisitions

Phase-contrast images were obtained by using the CKX53 system (Olympus, Tokyo, Japan), equipped with a charged-coupled device (CCD) camera (TrueChrome Metrics, Terratechnos, Tokyo, Japan). Confocal micrographs were captured using an FV3000 confocal laser-scanning microscope equipped with a uPlanSApo 40×/0.95 NA lens and analyzed with FV31S-SW imaging software (all from Olympus). Fluorescence intensity was measured by Zen Blue 2.6 software (Zeiss, Munich, Germany) with the background fluorescence subtracted. All images were further processed using Photoshop Elements 2018, according to the *Scientific Reports* guidelines.

### CARS microscopy measurements

The ultra-multiplex CARS microscope has been described in detail in a previous study [29]. Briefly, the fundamentals of the master laser source were used as the pump pulses. The wavelength, temporal duration, and repetition rate were 1064 nm, 50 ps, and 1 MHz, respectively. The near-infrared spectral component of the supercontinuum light source was used as the Stokes pulse. These beams were the outputs from the dual-fiber output and synchronized laser source (SM-1000, Leukos, Limoges, France). The pump power was adjusted using a variable neutral density filter. The two pulses were simultaneously introduced coaxially into an inverted microscope (Eclipse Ti-S custom-made, Nikon, Tokyo, Japan) via optical path adjustment and focused using a water-immersion objective lens (CFI Plan Apo IR 60X NA 1.27, Nikon). The sample was scanned using a piezo stage to reconstruct CARS images. Focal-plane imaging was performed at intervals of 0.5 μm/pixel. The CARS signal from the sample was collected using another microscope objective (S Plan Fluor ELWD 40x NA0.60, Nikon) and was detected using a spectrometer (LS-785, Princeton Instruments, Trenton, NJ, USA) and a CCD camera (PIXIS 100BR-UV-32, Princeton Instruments). The exposure time per pixel was adjusted in the range of 100–400 ms such that the CCD signal intensity was as strong as possible without saturating every pixel. The typical integration time was 180 ms. The Im[χ^(3)^] spectrum, which corresponds to the spontaneous Raman spectrum, was calculated using the maximum entropy method [60]. Baseline subtraction was performed using the following two-stage method. First, the baseline spectra outside the cells were averaged, and the spectrum was subtracted from the Im[χ^(3)^] spectrum of the cells. Next, we used the asymmetrically reweighted penalized least-squares method to subtract small variations in the baseline [61]. Bright-field images were obtained using a halogen lamp before and after the CARS microscope measurements.

### Spectroscopic analysis of CARS microscopy images

The obtained Im[χ^(3)^] spectra were fitted with the Levenberg–Marquardt nonlinear least-squares method using three Gaussian functions in the band (1540–1740 cm^-1^) that mainly contains the amide I mode owing to proteins [23,34]. The peak wavenumbers and linewidths of the three Gaussian functions are 1570 cm^-1^ and 15 cm^-1^, 1650 cm^-1^ and 30 cm^-1^, and 1667 cm^-1^ and 22 cm^-1^ (Fig. 1f) for (0) the purine ring skeleton of DNA and RNA (adenine and guanine) [29], (1) amide I of the protein α-helix structure and cis-type carbon double bonds of unsaturated fatty acids [29], and (2) amide I of the protein β-sheet structure, respectively [45,47,62,63]. The vibrational mode of water (1640 cm^-1^) had already been removed. The intensity of each Gaussian component at position 𝕏 is denoted as 𝑔_*n*, 𝕩_ (𝑛 = 0, 1, 2) when the position coordinates are 𝕏 = (𝑥, 𝑦, 𝑧). If the focus is on the nucleus, 𝑔_1, 𝕩_ mainly comprises a protein α-helix structure. We focused our analysis on two protein secondary structures, the α-helix and the β-sheet. The reason is that the spectral changes were caused mainly by the formation of unfolded/misfolded proteins and their aggregation. Amyloid-like aggregates induce extensive conformational changes from α-helices to β-sheets during their formation. Images were created for the ratios 𝑔_2, 𝕩_/(𝑔_1, 𝕩_ + 𝑔_2, 𝕩_).

For the images, we calculated the cross-sectional averaged values of the largest nucleolus in the nucleus and performed a comparison test between control and senescent cells (or proteasome-inhibited cells) as an index to distinguish senescent cells. If the values of the ratios were 10^-19^ or less, the positions in the images were displayed in dark purple. Similar to the fluorescent images of the indicator dyes, the subtraction 𝑔_2, 𝕩_ -𝑔_1, 𝕩_ was also performed to visualize the amount of amyloid-like aggregate. The subtraction contained the information of both the ratio and the amount concerning 𝑔_1, 𝕩_ and 𝑔_2, 𝕩_. Therefore, these values were related to the amount of β-sheet contained in the amyloid-like aggregates. The blue and red areas in the image corresponded to high (2) and high (1) components, respectively. The scale was based on the variance calculated for the values around the cell nucleus. This image analysis revealed relative amounts of the protein secondary structure.

### Statistical analysis

The results of phase-contrast and fluorescent image acquisitions are presented as the means ± standard deviation (SD) or as boxplots. Statistical significance was determined using a two-tailed t-test within Excel Software (Microsoft, Redmond, WA, USA). P < 0.05 was considered as a statistically significant difference.

The following are for CARS microscopy measurements. The normality of the data was tested using the Shapiro–Wilk method. Variance and comparison of DCB-treated cells and naturally binucleated ones were tested using Bartlett’s and Dunnett’s methods, respectively. Those of the MG132-treated cells were determined using an F-test and a Student’s *t*-test, respectively. The applications used for analysis were IGOR (ver. 8.0.4.2; WaveMetrics, Inc.), R functions (ver. 4.3.1), and R studio (2023.06.1). *P* < 0.05 was considered as a statistically significant difference. Each *P* value was indicated in the figure.

## Acknowledgments

This study was financially supported by Japan Society for the Promotion of Science KAKENHI Grant Number 18H02000 and 24K03251 (Grant-in-Aid for Scientific Research [B]), 21H04961 (Grant-in-Aid for Scientific Research [A]), and the Japan Science and Technology Agency Mirai Program grant (JPMJMI22G5) awarded to H.K.; KAKENHI Grant Number 19K06671 awarded to A.I. (Grant-in-Aid for Scientific Research [C]); and French government support managed by the National Research Agency under the Investments for the Future program (ANR-10-LABX-0074 Sigma-LIM) awarded to P.L. The authors gratefully acknowledge Juichiro Ukon, UKON CRAFT SCIENCE, Ltd., for assisting with the fruitful collaboration between the Japanese and French labs. The authors would like to thank Professor Toshiaki Hattori, Professor Kentaro Shiraki, and Dr. Hideaki Ito for their helpful discussions. The authors would also like to thank Kyosuke Tanaka, Dr. Shinichi Miyazaki, and Minori Masaki, for providing programs for analysis, and Zuliang Hu, Mia Ohbuchi, Sayuki Tokunaga, Minami Yoshimura, Yusuke Murakami, and Minako Suzuki for excellent technical assistance.

## Author contributions

S.I. developed the image analysis to detect unusual protein conformation. S.I., Y.O., H.K., and A.I. performed research: S.I., most of the spectroscopic experiments; A.I., biological experiments; Y.O., H.K., and A.I., priming the research. P.L. developed the laser source. S.I., A.I., and H. K. designed this cross-disciplinary study and wrote the manuscript. A.I. and H.K. supervised the study mainly about biology and spectroscopy, respectively.

## Competing interests

The authors declare no competing financial interests.

## Data availability

The datasets measured and analyzed during the current study are available from the corresponding authors upon reasonable request.

## Notes

### Competing Interest Statement

The authors have declared no competing interest.

### Summary of Updates

I have revised the title and abstract where it should be in Greek letters. No changes to the text.

